# Integration of public DNA methylation and expression networks via eQTMs improves prediction of functional gene–gene associations

**DOI:** 10.1101/2021.12.17.473125

**Authors:** Shuang Li, Cancan Qi, Patrick Deelen, Floranne Boulogne, Niek de Klein, BIOS Consortium, Gerard H. Koppelman, K. Joeri van der Velde, Lude Franke, Morris A. Swertz, Harm-Jan Westra

**Author notes:** These authors jointly supervised this work (Lude Franke, Morris A. Swertz, Harm-Jan Westra).

## Abstract

Gene co-expression networks can be used to infer functional relationships between genes, but they do not work well for all genes. We investigated whether DNA methylation can provide complementary information for such genes. We first carried out an eQTM meta-analysis of 3,574 gene expression and methylation samples from blood, brain and nasal epithelial brushed cells to identify links between methylated CpG sites and genes. This revealed 6,067 significant eQTM genes, and we observed that histone modification information is predictive of both eQTM direction and presence, enabling us to link many CpG sites to genes. We then generated a co-methylation network – MethylationNetwork – using 27,720 publicly available methylation profiles and integrated it with a public RNA-seq co-expression dataset of 31,499 samples. Here, we observed that MethylationNetwork can identify experimentally validated interacting pairs of genes that could not be identified in the RNA-seq datasets. We then developed a novel integration pipeline based on CCA and used the integrated methylation and gene networks to predict gene pairs reported in the STRING database. The integrated network showed significantly improved prediction performance compared to using a DNA co-methylation or a gene co-expression network alone. This is the first study to integrate data from two -omics layers from unmatched public samples across different tissues and diseases, and our results highlight the issues and potential of integrating public datasets from multiple molecular phenotypes. The eQTMs we identified can be used as an annotation resource for epigenome-wide association, and we believe that our integration pipeline can be used as a framework for future -omics integration analyses of public datasets.

We provide supporting materials and results, including the harmonized DNA methylation data from multiple tissues and diseases in https://data.harmjanwestra.nl/comethylation/, the discovered and predicted eQTMs, the corresponding CCA components and the trained prediction models in a Zenodo repository (https://zenodo.org/record/4666994). We provide notebooks to facilitate use of the proposed pipeline in a GitHub repository (https://github.com/molgenis/methylationnetwork).

## Introduction

Public resources like the Gene Expression Omnibus (GEO) databases have now accumulated data for tens of thousands of RNA-sequencing samples. The co-expression relationships derived from these data can be used to make inferences about the biological and molecular functions of genes^1^, which is particularly helpful for genes that currently lack biological annotation. To this end, we had previously created a gene co-expression network – GeneNetwork – of 31,499 RNA-seq samples from public resources and used it to provide biological annotations for genes and pathways^2^.

Similar to the RNA-seq data, a large amount of DNA methylation arrays for various types of tissues and diseases is now available in the public domain. This data is an invaluable resource for various applications, including biomarker discovery, and we may gain more mechanistic insights to identify novel functional gene–gene associations by analyzing this public RNA sequencing data together with DNA methylation data. However, we are not aware of any systematic attempts to use co-methylation to make inferences about gene function. To our knowledge there is currently no consensus resource that harmonizes the public methylation data and its corresponding contextual information, which limits re-use of the accumulated community resources. To overcome this, we harmonized and integrated 27,720 publicly available DNA methylation samples to create MethylationNetwork. Integrating MethylationNetwork with the already-existing GeneNetwork represents a powerful resource for genome annotation and functional research. However, to integrate the two networks, two major challenges needed to be addressed: we needed to identify links between genes and CpG sites and to integrate unmatched samples.

The first major challenge when harmonizing DNA methylation and RNA-sequencing datasets is that a gene is usually surrounded by many DNA methylation CpG sites. For most of these sites, it is still unclear whether changes in methylation status have molecular consequences and if they are correlated to gene expression changes. One way to determine this is to study the association between DNA methylation and gene expression levels (i.e. expression quantitative trait methylation; eQTMs^3,4^). Because most eQTM studies focus on a single tissue, it is often unclear whether the eQTMs identified in one tissue are consistently present in other tissues as well, and only a few eQTM studies have compared eQTMs across tissues^5^ or different cell types^6^. One previous study^5^, using 96 adult fetal liver samples and 85 adult subcutaneous and visceral adipose tissues, found that only up to 4% of eQTMs identified in one tissue could be found significant in another tissue; another study, using the umbilical cord of 195 newborn babies, observed that the overlapping ratio for eQTMs found in different cell types was between 0.13 and 0.42, and more than 85% overlapping eQTMs share the same effect direction. The tissue-specificity of eQTMs thus remains an open question. To investigate the tissue-specificity of eQTMs, we performed the largest-scale eQTM meta-analysis to date in 2,905 samples from blood, 473 samples from brain cortex and 196 samples from nasal brushed cells. This identified 33,877 significant *cis*-eQTMs (at a false discovery rate (FDR) < 0.05, reflecting 6,067 unique genes and 16,590 unique CpG sites). Out of these eQTMs, we found that more than 95% of overlapping eQTMs show the same effect direction. The detected and predicted eQTM results provide a valuable resource to study the downstream molecular effects of CpG sites associated with human phenotypes and diseases (epigenome-wide association studies; EWAS).

The second major challenge arose when trying to integrate publicly available DNA methylation datasets with publicly available RNA-seq datasets: the samples in public repositories are usually unmatched and have different disease and tissue compositions, whereas most available integration methods require that both methylation and RNA-seq data be available from the same individuals^7–9^. To resolve this, we first showed the complementary value of MethylationNetwork relative to GeneNetwork for the identification of known functional gene–gene associations. Furthermore, we developed an integration pipeline based on kernel cross-correlation matrix decomposition. Using this pipeline, we integrated GeneNetwork and MethylationNetwork and used the integrated results to predict functional gene–gene correlations that are collected in the STRING database and supported multiple lines of evidence. This integrated approach showed significantly better prediction results than either GeneNetwork or MethylationNetwork alone. Our integrated approach provides valuable insights into the process of harmonizing publicly available datasets containing unmatched DNA methylation and RNA-sequencing samples: insights that may need to be considered when integrating other publicly available datasets from different -omics layers.

We provide supporting materials and results, including the harmonized public DNA methylation data in https://data.harmjanwestra.nl/comethylation/, the discovered and predicted eQTMs, the corresponding CCA components, and the trained prediction models in a Zenodo repository (https://zenodo.org/deposit/4666994) and notebooks to facilitate use of the proposed pipeline in a GitHub repository (https://github.com/molgenis/methylationnetwork).

## Results

In this study, we first identified cross-tissue links between genes and CpG sites via an eQTM meta-analysis in blood, brain cortex and nasal brushed cells and then built machine learning models to predict eQTMs while using epigenetic data on histone modifications. We also collected and uniformly processed 27,720 public DNA methylation samples to build MethylationNetwork. With the discovered eQTMs and MethylationNetwork, we showed that MethylationNetwork could identify functional gene–gene associations that otherwise could not be seen in the corresponding co-expression patterns. Finally, to combine the advantages of both co-methylation and co-expression information, we developed an integration pipeline based on canonical correlation analysis (CCA) that integrates MethylationNetwork and GeneNetwork, and we illustrate the added value of the integrated network by showing its superior predictive power for known gene–gene associations compared to networks built solely from one source of information.

### Meta-analysis identifies eQTMs that are robust across different tissues

Genes are often surrounded by multiple DNA methylation CpG sites, which means that it is often unclear which CpG site should be linked to which gene. This information is, however, needed to integrate DNA methylation with RNA-sequencing datasets. Furthermore, EWAS studies have identified thousands of disease-associated CpG sites^10^, but the genes associated with most of these CpG sites are, which hampers functional interpretation. It also is often unclear whether such regulation would be tissue specific. To resolve these issues, we identified relationships between genes and DNA methylation CpG sites by performing an eQTM meta-analysis in blood, brain cortex and nasal brushed samples.

Our meta-analysis included 2,905 blood samples from the Biobank-based Integrative Omics Studies (BIOS) Consortium^4,11^, 473 brain prefrontal cortex samples^12^ and 196 nasal brushed samples, predominantly epithelial cells, from the PIAMA (Prevention and Incidence of Asthma and Mite Allergy) birth cohort^13^. In total, we tested 435,572 CpG sites and 25,488 genes and studied CpG–gene pairs that mapped within 1 mega-base pairs (mb) of CpG sites (Table 1, Supplementary Table 1). We discovered significant eQTM associations for 6,067 genes at an FDR of 0.05. The explained variance for gene expression levels was always lower than 0.4 (Supplementary Figure 1). We compared these associations to the largest eQTM analysis published to date by Bonder *et al.^4^* and a recent eQTM study in a child cohort by Ruiz-Arenas *et al*.^3^. As shown in Supplementary Figure 2, 83% (4,269) of the Bonder *et al.^4^* eQTMs were also found significant in our study, and 99% (4,246) of eQTMs were of the same direction. Only 16% (3,861 out of 23,862 eQTMs) of the Ruiz-Arenas *et al.^3^* eQTMs were also significant in our study, but 97% (3,731 of the 3,861) had consistent effect direction. The small overlap of eQTMs between our study and Ruiz-Arenas *et al.^3^* could reflect differences between adult and child cohorts, as Ruiz-Arenas *et al*. also observed.^3^ We further identified eQTM associations for 2,243 additional genes compared to Bonder *et al*.^4^ and 5,872 additional genes compared to Ruiz-Arenas *et al*.^3^ As expected, the majority of the eQTM genes (n=4,176; 67%) were negatively correlated with the nearby CpG sites (Table 1). We tested whether this directionality was consistent among different tissues by replicating the discovered eQTMs in each of the three tissues separately and observed that over 98% of the eQTMs identified significantly in both the meta-analysis and in each tissue shared the same direction (Figure 1). This high concordance was also observed when comparing between datasets (Supplementary Figure 3). The summary statistics of the eQTMs discovered in each of the cohorts are provided in Supplementary Tables 2–4. We therefore conclude that the eQTMs identified by our meta-analysis are highly reproducible in different tissues, with only a few eQTMs exhibiting significant tissue-specific effect directions. We note that only a small portion of the significant eQTMs from the meta-analysis could also be detected as a significant result when using only samples from the three tissue datasets: 5,398 of the 6,067 eQTM genes could be detected when using only the BIOS dataset, 124 were detected when using only the ROSMAP dataset and 69 when using only the PIAMA dataset. Additionally, among significant eQTMs with consistent effect direction, the effect sizes were generally different (Figure 1), which could be due to differences in sample size. For the subsequent analysis, we used the eQTMs with consistent effect direction as an empirical strategy to link CpG sites to genes that can be used in multiple tissues.

**Figure 1:**
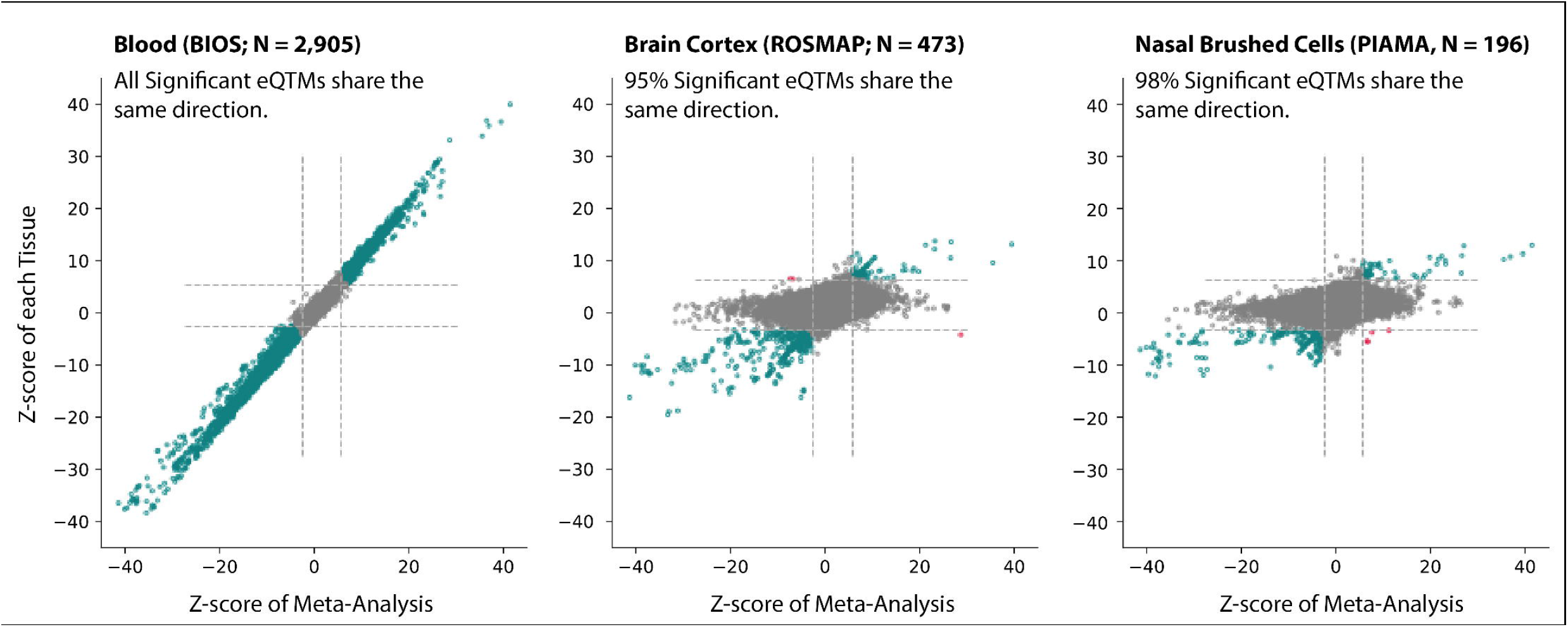
Meta-analysis identified eQTMs with consistent directions across blood (BIOS), brain cortex (ROSMAP) and nasal brush (PIAMA). Each dot represents an eQTM. The Z-score of its association coefficient from the meta-analysis is indicated on the x-axis, and that from the single-tissue analysis is indicated on the y-axis. Dashed lines indicate the significance threshold (FDR < 0.05) calculated from the meta-analysis and from each corresponding tissue. Gray dots indicate that the eQTM is not significant in the meta- or single-tissue analysis. Pink dots indicate that the eQTM is significant in both analyses and show opposite directions. Green dots indicate that the eQTM is significant in both analyses with consistent direction.

**Table 1.** Summary of the eQTM analysis.

|  | BIOS (Blood) | ROSMAP (Brain Cortex) | PIAMA (Nasal brushed Cells) | Meta Analysis |
| --- | --- | --- | --- | --- |
| Individuals | 2905 | 473 | 196 | 3574 |
| Tested genes | 19942 | 19144 | 16859 | 25488 |
| Tested CpGs | 402016 | 420963 | 415538 | 437572 |
| significant eQTMs | 39002 | 988 | 688 | 33877 |
| Unique eQTM genes | 6628 | 197 | 202 | 6067 |
| Unique positive eQTMs genes | 2181 | 48 | 29 | 1891 |
| Unique negative eQTM genes | 4447 | 149 | 173 | 4176 |

We next examined the negative and positive eQTMs for their different properties. Specifically, we examined the distance between the CpG site and the transcription start site (TSS) of the eQTM genes, and whether the CpG sites co-localize with specific histone modifications identified in ROADMAP^14^ (Methods). Negative eQTM CpG sites are significantly more likely to be located more proximally to the TSS of a gene (Figure 2, Chi-square test two-sided p-value: 1.0×10^-65^), with 60% of negative eQTM CpGs located within 50kb of the TSS versus 32% of positive eQTMs. We had previously shown that histone modifications are informative for predicting the direction of eQTMs (Bonder et al, Nature Genetics 2017), and we used the same method here to evaluate the overlap ratio between histone modification data and CpG sites across different cell types in the ROADMAP epigenomics project^14^. Specifically, the overlap ratio represents the proportion of cell types in which a CpG site overlaps with a histone modification. Similar to the observations in our earlier publication^4^, we found that the overlap between the CpG sites from the eQTMs and histone modifications from ROADMAP^14^ (Methods) indicates a complex relationship between TSS distance and eQTM directions. Figure 2 shows an example of this in which the overlap with H3K27ac is significantly higher (Mann-Whitney U rank test two-sided p-value: 3.2×10^-32^) for negative eQTMs than for positive eQTMs. However, when the TSS distance increases, the overlap ratio with H3K27ac with negative eQTMs gradually decreases while it increases for positive eQTMs.

**Figure 2:**
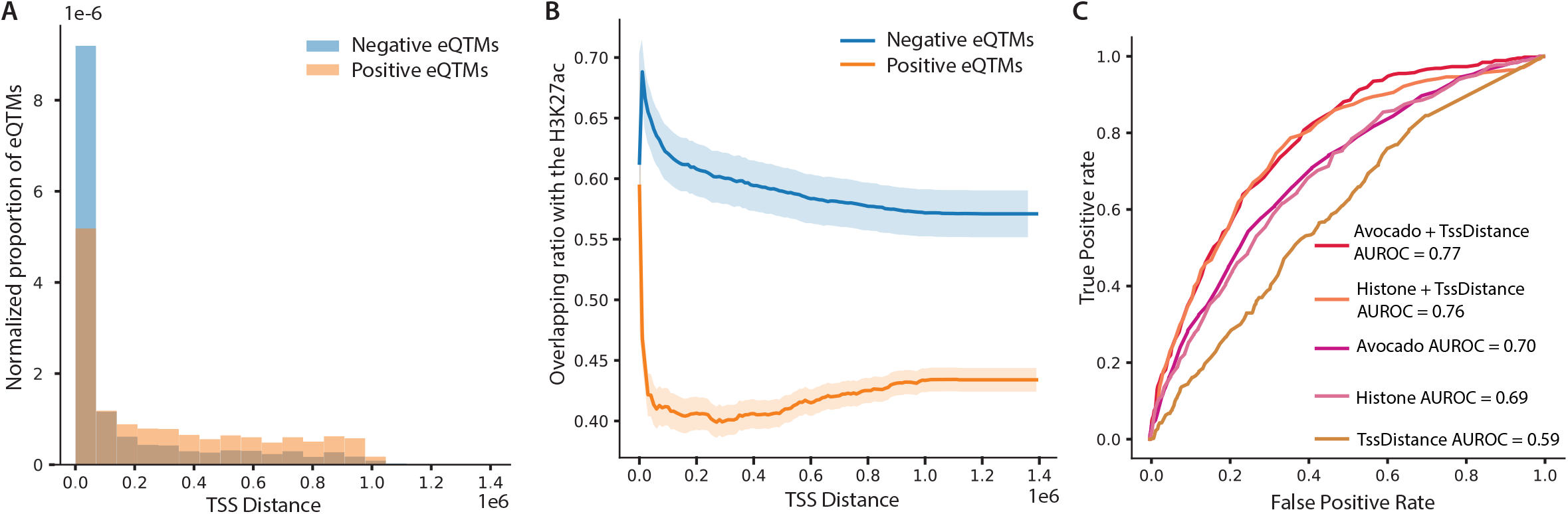
Depending on transcription start site (TSS) distance, positive and negative eQTMs show different overlapping properties with histone modifications. **a** TSS distance distribution for negative and positive eQTMs. y-axis indicates the normalized eQTM counts. The normalized counts are calculated so that the total area of the histogram adds up to 1. **b** Overlapping ratio between CpG sites with the histone modification H3K27ac with regards to TSS distance for negative and positive eQTMs. **c** Performance of the models that predict eQTM direction using the ROC and AUROC value. Legend indicates the feature used in each model.

Based on this observation, we constructed a machine learning model to predict the direction of eQTMs using histone modification data from the ROADMAP project^14^. We had previously attempted to predict these directions^4^, but here we improved these predictions in several ways. First, while the previous study used a single decision tree for this classification problem, we used random forest algorithm, which aggregates results from multiple deicion trees, lowering the prediction variance, and therefore increasing the overall predictive power. Second, we tested different representations of the histone modification data: 1) we used the representation used by the previous study, which is the ratio of overlapping histone modifications among different cell types, and 2) we used the representation from the Avocado study^15^, in which the histone modification data is summarized by matrix decomposition. Beyond the changes in model training, we adopted a more stringent evaluation criterium that assesses only one CpG per gene (previously multiple CpGs were assessed per gene). Here, we observed that the Avocado representation had the best performance when modeled together with the TSS distance, resulting in an area under the receiver operating characteristic (AUROC) of 0.77 (Figure 2). Furthermore, using a separate random forest-based model, we predicted whether a CpG site is associated with a gene (i.e. the existence of an eQTM) using histone overlap features, and this model achieved an AUROC of 0.94 (Supplementary Figure 4, Supplementary Figure 5; Supplementary Table 5). These models indicate that relationships between DNA methylation and gene expression levels can be reliably predicted using the distance to the TSS and histone modifications.

### MethylationNetwork, constructed with 27,720 public DNA methylation samples, captures tissue differences

Using the 6,067 eQTMs identified from our eQTM meta-analysis, we proceeded to link these two -omics data levels from public repositories. We decided to use these eQTMs instead of the ones predicted from the random forest models, to provide any influences from potential false positives. We processed raw data from 27,720 Illumina 450k methylation arrays collected from 532 projects previously deposited in the GEO, performed DASEN normalization^4^ and created MethylationNetwork after outlier removal (Supplementary Figure 6, Methods). We also harmonized the accompanying sample annotations by defining 32 tissue groups and 90 disease groups, with each group comprised of more than 10 samples (Figure 3, Supplementary Table 6, Supplementary Table 7). We then performed principal component analysis (PCA) on the methylation data, represented the data using UMAP^16^ and observed that the samples in our methylation network clustered according to tissue group rather than per study (Figure 3). This suggests that our harmonization and DNA methylation normalization steps removed potential batch effects, even though our dataset contained samples from 532 projects. Furthermore, this clear tissue clustering reassured us that MethylationNetwork carries biological meaning even with the potential batch effects introduced by various experimental protocols.

**Figure 3:**
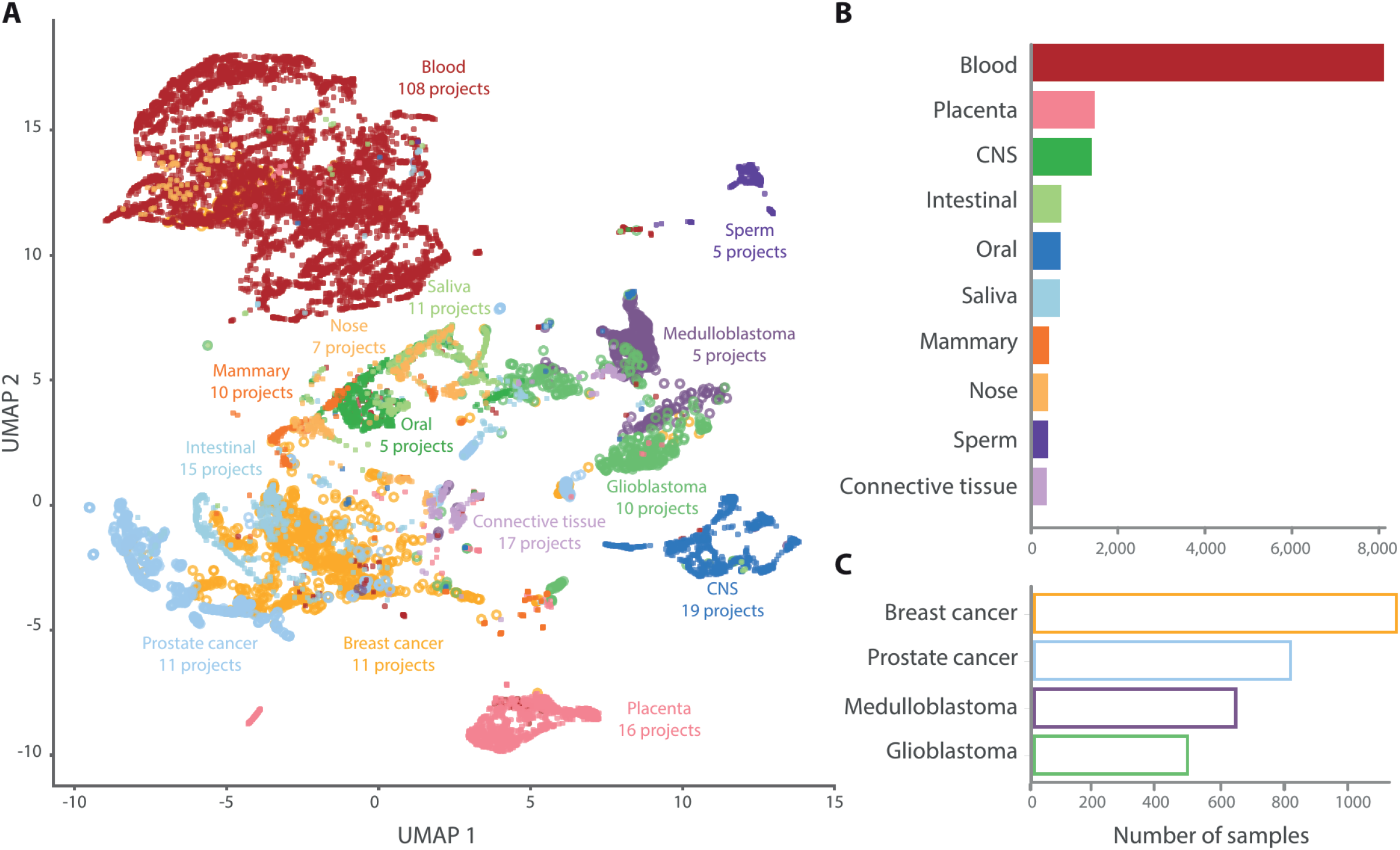
Overview of public methylation data from GEO databases. **a** UMAP plot of the first four PCA components from the public methylation data. Each sample is colored by the tissue origins. Next to the tissue labels, the number of involved projects with unique GSE number is also indicated (e.g. the blood samples are collected from 113 projects). **b** the number of samples from each non-cancer tissue. c the number of samples from each brain cancer. For better visualization, we show only the 10 most abundant non-cancer tissues and the 4 most abundant brain cancer tissues. Annotations for each sample can be found in Supplementary Table 6.Full description of tissue composition can be found in Supplementary Table 7.

Previously, many researchers have examined the TCGA database and shown that DNA methylation data is informative for discriminating between cancer types and identifying the primary tissue of origin. We can now recapitulate these results using many different publicly available cancer studies: we trained a Support Vector Machine (SVM) model to predict tissues using PCA components from MethylationNetwork, identified potential DNA methylation biomarkers for each tissue (Supplementary Figure 7, Supplementary Table 8) and applied the prediction model to predict the origin tissues of metastasis samples. For the metastasis origin tissue prediction, we used four projects that encompass data from 161 metastases that occurred in a tissue different than the primary tissue of origin. These included metastases in brain (GSE108576), prostate (GSE38240) and colorectal tissue (GSE75546 and GSE77954). For 80% of the metastasis samples, the model predicted the correct tissue of origin as the top prediction. For 90% of the samples, the model listed the correct tissue of origin in the top three predictions. For 95% of the samples, the correct tissue was listed in the top five predictions (Supplementary Table 9, Supplementary Table 10). As such, our procedure and harmonized DNA methylation dataset can be useful to determine tissue of origin for a metastasis sample.

### MethylationNetwork identifies known functional gene–gene associations not seen in GeneNetwork

By applying MethylationNetwork and using eQTMs as bridges between the CpG sites in MethylationNetwork and the genes in GeneNetwork, we next compared the co-methylation patterns between eQTM gene pairs to their co-expression counterparts from our previously published gene co-expression network consisting of 31,499 RNA-seq samples^2^. We examined whether co-methylation provides complementary information for identifying the functional or physical associations between genes. We collected the gene pairs that consistently showed high co-methylation but low co-expression and checked whether those gene pairs were enriched for functional or physical associations known in the STRING database^17^.

Our analysis confirmed that co-methylation relationships are a complementary resource to co-expression relationships for identifying true functional or physical gene–gene associations. As shown in Figure 4 and Supplementary Figure 8, the correlations between co-methylation and co-expression were low, with a Pearson correlation coefficient of 0.02 for the BIOS dataset and 0.03 for the MethylationNetwork and GeneNetwork. To determine which genes showed distinct patterns between methylation and gene expression data, we selected gene pairs with a high absolute co-methylation value (Spearman r > 0.7 or r < −0.7) but a low co-expression value (Spearman r < 0.5 and r > −0.5). This identified 4,001 gene pairs, including 168 pairs that were also reported in the STRING database. This ratio (171/4001=0.04) was significantly higher than random (chi-square test p=0.001), indicating that co-methylation patterns are enriched for known gene–gene associations. Specifically, 6 out of the 171 gene pairs (Supplementary Table 11) were directly supported by evidence annotated as “experiments” in the STRING database but not by evidence annotated as “co-expression”. This included the *MRPL35-MRPS18B* gene pair, which we observed to be highly co-methylated in both the BIOS dataset (Spearman r = 0.86) and MethylationNetwork (Spearman r = 0.71) but we observed no co-expression in either dataset (Spearman r = −0.10 in BIOS and r = 0.28 in GeneNetwork; Figure 4 and Supplementary Figure 9). For the remainder of the 165 gene pairs, 14 were supported by “databases” evidence and 109 by “textmining” evidence (Supplementary Table 11).

**Figure 4:**
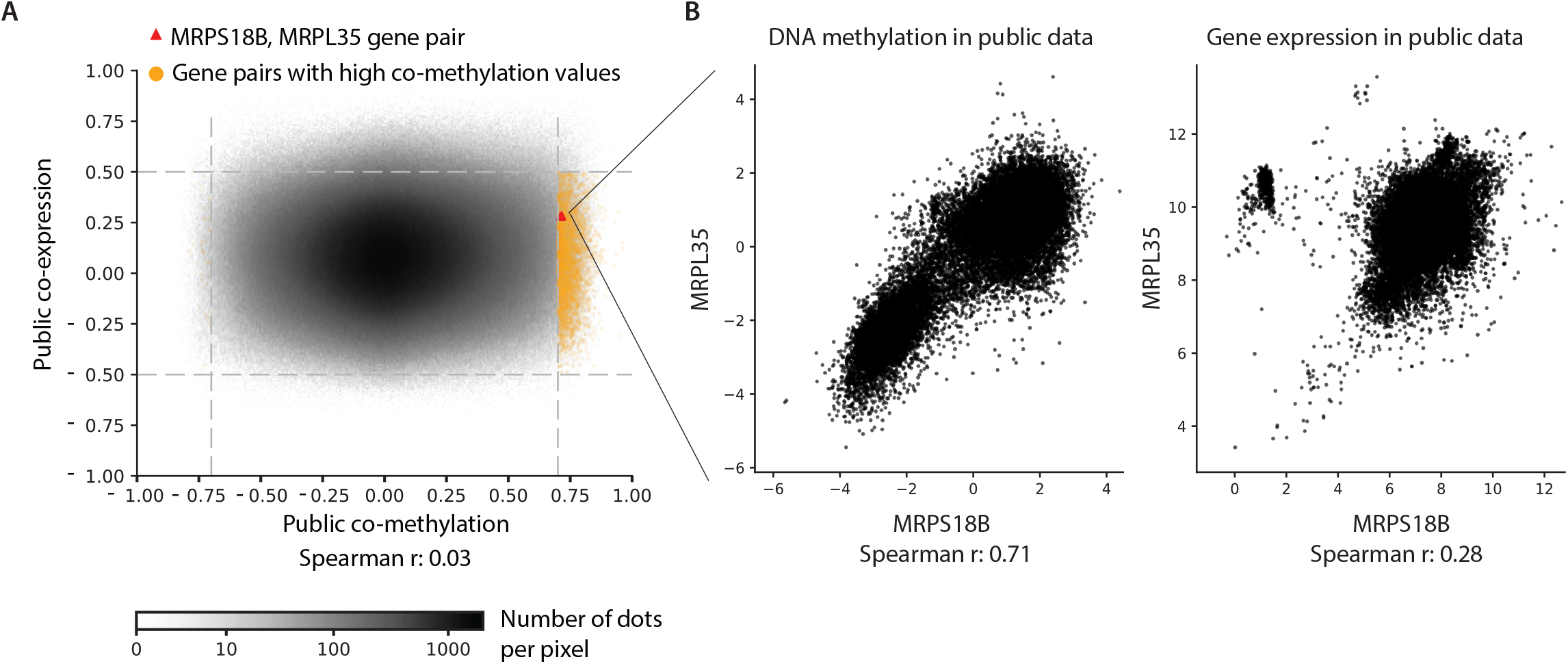
Comparison between Co-expression and Co-methylation patterns in the public datasets. **a** density plot showing the comparison for co-expression and co-methylation values for any gene pairs from the eQTM genes in the public data. Yellow dots indicate the 3,974 gene pairs with high co-methylation values (r > 0.7) and low co-expression values (r < 0.5). The red triangle indicates the *MRPL35-MRPS18B* gene pair that is highlighted in **b**, which show the DNA methylation and gene expression levels in the public datasets for these two genes.

### Co-methylation patterns identify functional gene–gene relationships that are complementary to gene-co-expression patterns

Since we observed that co-methylation data can provide complementary information to co-expression relationships and that the co-expression and co-methylation are only weakly linked, we next evaluated whether we could put both co-expression and co-methylation directly into a linear model to predict known gene–gene pairs from the STRING database. We compared this prediction performance against a baseline model that only uses co-expression data. However, this approach did not show a significant prediction improvement when combining the two networks (Figure 5D, E). This could partly be due to the large difference we observed between co-methylation and co-expression patterns for the same gene pairs (Supplementary Figure 8). We reasoned that, because of the complex interplay between gene expression and DNA methylation levels, this method of concatenation was not able to use the complementary values from these two -omics layers.

**Figure 5:**
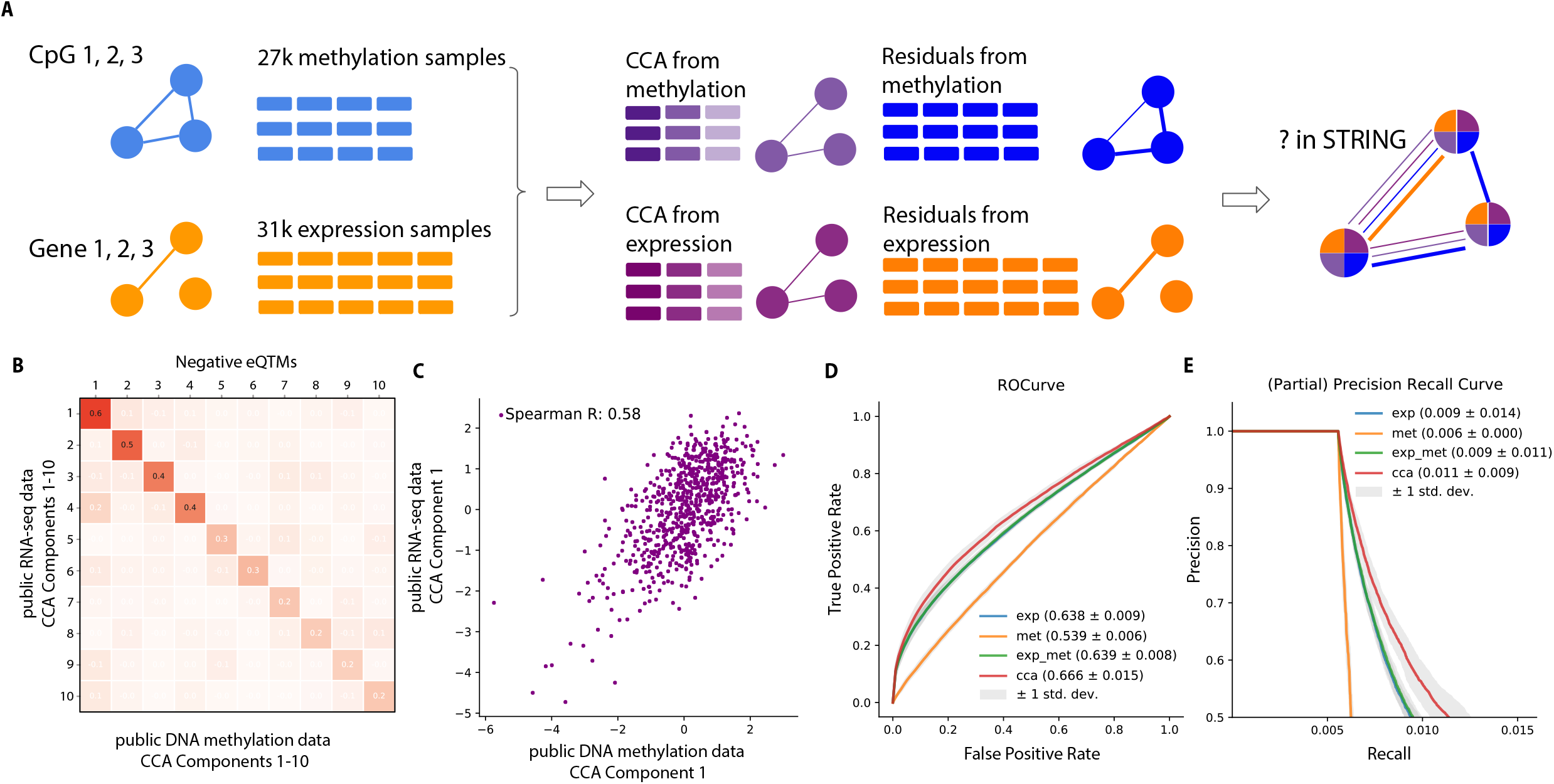
CCA components predict gene interactions significantly better than co-expression alone. **a** Flowchart for the CCA-based prediction pipeline. **b** Correlation of CCA components obtained from the public RNA-seq data and DNA methylation data. **c** Scatter plot showing the correlation between the first CCA component obtained from the public RNA-seq data and DNA methylation data. **d** ROC curves showing the performance comparison for the prediction of gene-gene associations **e** Partial precision and recall curves showing the performance comparison for the prediction of gene-gene associations. For both (**d**) and (**e**), uncertainty is quantified through 5-fold cross validation. The “exp” indicates the model constructed with only co-expression values. The “met” indicates the model constructed with only co-methylation values. The “exp_met” indicates the model constructed with both co-expression and co-methylation values. The “cca” indicates the model constructed with the CCA components and residuals from the pipeline indicated in **a**.

To better leverage the complementary information between GeneNetwork and MethylationNetwork, we developed an integration pipeline based on canonical correlation analysis (CCA; Figure 5A) and eQTM associations. The first step in this pipeline consists of identifying CCA components using the methylation and gene expression values, using eQTMs as links between genes and CpG sites. We evaluated the performance of this first step by examining the correlation of CCA components and the intermediate results (PCA components, see Methods, Figure 5B, C, Supplementary Figures 10–12). We also explored the gene enrichment results for the CCA components (Methods, Supplementary Table 12).

Here, we observed that the best performance from the CCA pipeline was obtained by analyzing negative and positive eQTMs separately (Supplementary Figure 12). Analyzing only negative eQTMs returned four components that showed a Spearman correlation that passed the nominal significance level of 0.05, while analyzing the entire eQTM dataset returned fewer components with lower correlations in general. The CCA analysis with positive eQTMs returned the lowest correlations, likely because of the small number of available positive eQTMs. Moreover, we also observed that the CCA components from the separate analysis of positive and negative eQTMs showed distinct enrichments for gene ontology terms (Supplementary Table 12). Based on this observation, we analyzed the negative and positive eQTMs separately in the following analysis. The variance explained by the CCA components was generally low (Supplementary Figure 13). For example, the average explained variance for CCA component 1 from the negative eQTMs in MethylationNetwork was 0.012.

The second step of our pipeline aims to predict functional gene associations defined in the STRING database. To do so, we first determined the CCA components, which captures high correlated components between gene expression and DNA methylation data. Additionally, we determined the components that are specific to gene expression and methylation data by determining the residual gene expression and methylation levels after correcting for the CCA components using linear regression. For both these residuals and the CCA components, we then calculated a pair of correlation matrices: one for the gene expression values and one for the methylation values. Finally, we used the resulting correlation matrices to describe each eQTM gene pair and used these correlations to determine the probability that each gene pair had a known functional association in the STRING database using a logistic regression model. We defined gene pairs with a known functional association by setting two thresholds (400 and 600) on the “combined_score” provided in the STRING database.

We observed that our integrative approach significantly improved the prediction results compared to MethylationNetwork and GeneNetwork alone. We compared area under the receiver operating and precision recall curves (AUROC and AUPRC) values for the following four approaches: 1) co-expression values from the GeneNetwork, 2) co-methylation from the MethylationNetwork, 3) co-expression and co-methylation values both networks and 4) the CCA components and the residuals from both networks. We quantified the uncertainty with 5-fold cross validation. As shown in Figure 4D, the fourth approach using the integrated CCA pipeline yielded an AUROC of 0.67, which was significantly higher than the AUROC from the first (unpaired two-sided t-test p-value: 0.004) and third approach (unpaired two-sided t-test p-value: 0.005).

We observed that the CCA approach did not show significantly higher AUPRC values than the first approach. However, it was significantly better at identifying true positive gene pairs. When focusing on predictions with a precision > 0.5, we observed a significantly higher (two-sided t-test p-value: 0.004) average recall value for the CCA approach (0.011) compared to using only GeneNetwork (0.009; Figure 5E). Furthermore, the CCA approach also showed significant better AUPRC values when we adopted a more stringent threshold (“combined_score” > 600) for the STRING gene pairs (Supplementary Figure 14).

The regulation of gene expression levels may be dependent on post-translational modifications (PTM). We previously suggested that such PTMs may prevent a gene network from fully recovering gene–gene relationships. We therefore investigated whether our CCA approach could recover these relationships by better predicting gene pairs undergoing PTM. To do this, we compared the prediction scores for PTM gene pairs with those for non-PTM gene pairs. We defined PTM gene pairs using genes that are annotated with PTM and at least one gene–gene interaction in the Uniprot database^18^. We observed that the CCA approach scored the PTM gene pairs significantly higher (two-sided t-test p-value: 1.46×10^-27^; Supplementary Figure 15). Furthermore, the CCA approach scored these gene pairs significantly higher than any of the other approaches, while the scores for non-PTM gene pairs were not significantly different (Supplementary Figure 15).

We also explored another application of our integration pipeline by testing whether we could use the integrated public datasets to provide evidence for new eQTMs. We first tested whether the distributions of the eQTMs discovered in the meta-analysis were significantly different from a random pair set of CpG sites and genes. To do this, we correlated the CCA components from the public datasets for the eQTM pairs and obtained the correlation distribution. We then performed the same steps for a random set of CpG sites and genes and obtained the background distribution. We observed that the correlation distribution for eQTMs was significantly different from the random set of CpG sites and genes (Supplementary Figure 16). We next tested whether the predicted eQTMs from our previous results also share the same property as the meta-analysis eQTMs and observed that the predicted eQTMs also displayed a significantly different distribution from a random set (Supplementary Figure 17).

In conclusion, we have shown that MethylationNetwork provides complementary value to GeneNetwork for identifying known gene–gene associations. Although both networks show very few common properties, the CCA integration pipeline identified common components enriched for common Gene Ontology (GO) terms for eQTMs. We show that this integrative approach can predict known gene–gene associations significantly better than GeneNetwork alone.

## Discussion

Methylation profiles are highly tissue specific. However, it is unknown whether the correlations of CpG sites with nearby genes – eQTMs – are also tissue-specific, and only a few studies have investigated the tissue-specificity of eQTMs. One previous study discovered eQTMs using liver tissues from 96 adult individuals and muscle and adipose tissues from 85 individuals^5^. They intersected the significant findings and discovered that only 4% of the significant results were consistent across tissues. Another study concluded that the directions of eQTMs are mostly consistent in different blood cells^6^. In our study, in which we recruited larger datasets and tested in three different tissues, we observed consistency for the effect direction of eQTMs. However, we also observed that many significant eQTMs in the meta-analysis cannot be replicated as significant in another tissue. Even for those eQTMs that can be replicated significantly, the effect sizes can be quite different in different tissues. This could be partly explained by the relatively smaller sizes of the brain cortex and nasal brushed cell datasets used in our meta-analysis. Therefore, the tissue-specificity of eQTMs requires further investigation through the collection of larger datasets of comparable size from various tissues. Moreover, large, paired datasets from non-blood tissues are needed to directly answer the question of tissue-specificity of eQTMs.

The directions of eQTMs were important for improving the performance of the integration pipeline. We suspect that the eQTMs with different effect directions reflect different epigenetic regulatory mechanisms, based on the complex association changes we observed between the location of eQTM CpG sites and various histone modifications in terms of varying TSS distance. Furthermore, we hypothesize that these two types of eQTMs maybe involved in different biological pathways because we observe that the CCA components obtained when analyzing negative eQTMs and positive eQTMs separately were enriched for distinct GO and Human Phenome Ontology (HPO) terms. We therefore argue that the negative and positive eQTMs cause different changes in expression and DNA methylation levels. As a result, we chose the simplest solution, i.e. analyzing the negative and positive eQTMs separately, and indeed observed that the CCA components from MethylationNetwork and GeneNetwork correlate better when the negative and positive eQTMs are analyzed separately than when they were analyzed together (Supplementary Figure 11). These observations underscore the importance of using the underlying biological mechanisms when designing statistical pipelines for -omics integration studies.

In our eQTM study, we evaluated single CpG sites that influence nearby gene expression. However, it is conceivable that multiple CpG sites might independently influence gene expression levels of one gene. Consequently, the eQTMs we identified might not capture the full variation of gene expression regulated by methylation. One solution to this is to use multiple linear regression, a framework comparable to Predixcan^19^. This would also solve the many-to-one relationships currently present in eQTM datasets. Finally, distal effects from CpG sites on gene expression levels could also play an important role, so a statistical framework that allows integration of *cis-* and *trans*-eQTM effects could further increase the gene expression variance explained by eQTMs.

## Conclusion

In this study, we discovered and predicted *cis-*eQTMs from a meta-analysis using blood, brain cortex and nasal brushed cells. We harmonized 27,720 DNA methylation samples from public resources to create MethylationNetwork, which showed that DNA methylation data is highly variable between tissues. With the identified eQTMs, we then developed a CCA-based pipeline to integrate MethylationNetwork with GeneNetwork, a previously created gene co-expression network. Using the integrated datasets, we saw significant improvements in the prediction of known gene–gene associations reported in STRING database. Our work provides the largest meta-analysis for *cis*-eQTM discovery, is the first large-scale examination of the tissue-specificity of eQTMs and the first to harmonize public DNA methylation data across different tissues and diseases. Finally, we present a novel integration pipeline for unmatched public datasets. We believe that our work facilitates the re-use of existing public data, makes the case for large-scale eQTM analysis across tissues and for result interpretation with other epigenome assays, and highlights the importance in incorporating biological mechanisms for multi-omics integration methods.

## Methods

### Meta-analysis for eQTMs in blood, brain cortex, and nasal brushed cells

To perform meta-analysis across different tissues and identify the potential regulatory effects of CpG sites on the nearby genes (eQTMs), we first used 2,905 blood samples from the BIOS consortium^4,11^, 196 nasal epithelium samples from PIAMA cohort^13^ and 473 brain cortex samples from the ROSMAP study^12^. For the DNA methylation data from each cohort, we used the M values and performed the following steps for normalization: quantile normalization, centering for each probe and standardization of the values for each sample. Next, we corrected each of these datasets for potential batch effects separately by regressing out principal components determined over the sample correlation matrix. We then regressed out the first 5, 10 and 22 principal components from PIAMA, ROSMAP and BIOS data, respectively. We further removed variance in the expression data that might not be attributed to eQTMs by regressing out genetic effects of nearby SNPs (i.e. *cis*-eQTLs and *cis*-meQTLs). We repeated this process iteratively until no more significant *cis*-eQTLs or *cis*-meQTLs were detected. Then, we mapped *cis*-eQTMs using Spearman’s rank correlation, testing CpG sites located within 1Mb of the TSS of each gene. To correct for multiple testing, we performed 10 permutations to empirically control the probe-level FDR rate at 5%, a procedure similar to previous work^20^. To examine whether the eQTMs identified by our meta-analysis share the same direction in different tissues, we replicated the 33,877 significant eQTMs in each of the three tissue datasets separately.

### Prediction of eQTMs

To predict potential eQTMs that were not detected by our meta-analysis, we created a machine learning–between the methylation site to the TSS distance.

To represent the histone modification information, we compared two different feature representation methods: the Avocado representation^15^ and a representation used in a previously published eQTM study^4^. Briefly, the Avocado representation^15^ was created by factorizing the epigenome data matrix from the Roadmap^14^ and FANTOM5^21^ consortia and learned latent factors that represent the cell and assay types and the genomic axis. We used the latent factors along the genomic axis as the input features for the eQTM prediction models. The previously published study^4^ used the colocalization ratio between a CpG site and histone modification peaks among the 127 cell types measured in the Roadmap consortium. Specifically, when a CpG site co-localized with a specific histone modification peak for all cell types, we used an input feature of 1, and when the CpG site only co-localized with the histone modification peak among 50% of the cell types, we used an input feature of 50% for this specific CpG site and histone modification combination.

Using the histone modification features and TSS Distance, we built the two machine learning models. The first model predicts whether there is a *cis*-eQTM association (or not) for any CpG site and its nearby gene (within 1Mb). We used a threshold of 0.8 for the confidence score from the first model to determine whether a CpG–gene combination was likely an eQTM. We determined this threshold such that the precision from the test dataset was 90%. The second model predicts whether the association is negative or positive for the recognized *cis*-eQTMs from the previous model.

For the prediction pipeline, we constructed the training and test datasets as follows. For the eQTM prediction model, we first annotated the 33,877 significant eQTMs discovered in the meta-analysis with histone modification features and TSS Distance. We then deleted all eQTMs with missing feature values, which resulted in 33,866 eQTMs, and randomly selected an equal number of non-eQTM CpG–gene pairs to form a balanced dataset. We randomly split the eQTMs into training (75% of data; 25,347 eQTMs and 25,452 non-eQTM CpG–gene pairs) and test (25% of the data; 8,519 eQTMs and 8,414 non-eQTM CpG–gene pairs) datasets. For the model to predict the eQTM direction, we first annotated all the eQTMs with histone modification features and TSS Distance, then selected unique CpG sites that are potentially regulating unique genes in our eQTM CpG–gene set. If one CpG site was associated with more than one gene, we selected the eQTM with the lowest p-value. This resulted in 4,624 eQTMs. We selected the eQTMs with an FDR between 0.05 and 0 as the training dataset and the eQTMs with an FDR below 0.01 for testing.

To gain a credible set of predicted eQTMs, we used a probability score threshold of 0.99 for the eQTM prediction model and assumed that the resulting eQTMs are likely eQTMs. The threshold 0.99 was chosen so that the precision value for the test dataset of this model achieves 98%. The corresponding recall value was 26%. We also trimmed the results from this model to have at most 28 genes associated with a CpG site. The trimming gene limit was the maximum number of genes that were associated with one CpG site in the eQTMs detected by our meta-analysis. For the predicted eQTMs, we further applied the eQTM direction prediction and used a probability score threshold of 0.4 to determine the positive eQTMs and a probability score threshold of 0.3 to determine the negative eQTMs. We deemed eQTMs with a probability higher than 0.4 to be a positive eQTM, and those with a probability value below 0.3 to be negative eQTMs. The threshold for positive eQTM was chosen so that the recall was reasonably high (54%), and the precision value was 45%. The threshold for negative eQTMs was chosen so that the precision was reasonably high (94%), and the recall value was 50%.

### Public methylation data preprocessing and normalization

To harmonize the methylation data, we collected all available public Illumina Infinium 450k Idat files from the GEO database^22^ (dated 19-03-2020). For each project with a unique GSE number, we preprocessed it with the DASEN pipeline^4^ and then applied quantile normalization. Later, we concatenated all normalized datasets and proceeded with normalization steps. First, we used PCA for outlier detection. For all samples, we obtained the values for the first principal component, calculated a Z-score per sample and discarded the samples with a Z-score > 3 (Supplementary Figure 6). After outlier detection, we performed the following normalization steps: quantile normalization, centering values for each CpG site and Z-transformation for each sample. Using these steps, we obtained a uniformly processed methylation dataset with 422,803 CpG sites for 27,720 samples. Additionally, we collected the sample annotations from the GEO dataset using the E-utilities API^23^ and manually curated the tissue and disease labels. The harmonized annotations are provided in Supplementary Table 6.

### Tissue prediction and biomarker discovery

To identify potential biomarkers for tissues and diseases using the public methylation dataset, we first standardized the processed public methylation dataset. We then performed PCA over the CpG site correlation matrix and used the principal components to describe each sample and the eigen vectors to describe each CpG site. With the first 100 principal components representing the samples, we trained support vector machines with linear kernels for tissue predictions. For the tissue prediction model, we selected 24 tissues from the 35 tissue groups collected. This selection was based on two criteria: 1) samples were collected from at least two projects with unique GSE numbers, so that we could avoid overfitting the tissue prediction model to the batch effects introduced by, for example, use of different protocols when generating the data, and 2) the sample size was larger than 100, so that there were enough data points to train and validate the tissue prediction model.

For both the tissue and disease prediction models, we applied SVM with a linear kernel as the multi-classification engine and standardization for feature preprocessing and used accuracy, recall and precision together with the F1-score as the evaluation matrix. The model construction steps were carried out with scikit-learn Python package^24^, and the scripts can be found in our GitHub repository (see Web resources). As an independent evaluation dataset, we also tested the tissue prediction model on four metastasis projects (GSE108576, GSE38240, GSE75546 and GSE77954). The metastasis projects contained 60 intestinal samples, 37 skin samples, 30 mammary samples, 18 lung samples and 12 gonadal samples. We explicitly removed these samples from our normalized public dataset, so they were not seen by the model during the model construction process. We applied the trained model on the metastasis datasets and took the top three tissue prediction results to compare with the actual tissue origin to determine how many origin tissues were correctly predicted as the first prediction or among the top three predictions of the model. The predictions and their true tissue origins are provided in Supplementary Table 11. We also provide the principal components of the public methylation dataset and the trained models in a Zenodo repository (see Web resources).

To identify potential CpG site biomarkers for each tissue, we calculated the feature importance as a proxy for the likelihood that a CpG site is a biomarker. The higher the feature importance, the more the value of a CpG site contributes to the tissue prediction results and the more likely it is that this CpG site could be used as a potential biomarker for that tissue.

Because we used PCA transformation beforehand as a low-dimensional representation for the CpG sites and then used the eigenvectors for each CpG site as the input for the tissue prediction model, we calculated the feature importance for each CpG site by combining the coefficients from both the PCA transformation and the SVM model.

Formally, we denoted the eigen vector index with *n* and the total number of eigenvectors as *N* as the input for our SVM model. We denote the tissue index with *t* and the total number of tissues tested *T* in our tissue prediction models. As such, the feature importance of a CpG site in predicting tissue *t* is given by the following Equation 1:

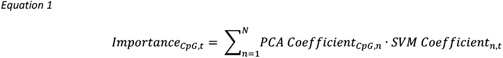

To evaluate the identified CpG biomarkers for tissues by the feature importance, we calculated Z-scores for each tissue and selected those CpG sites with a Z-score > 3 (i.e. those with a feature importance value outside the 3 standard deviation range from the average value). We then examined whether their methylation values were significantly different in the corresponding tissue when compared to CpG sites that are important for other tissues. To do this, we used the Kolmogorov-Smirnov test to test whether there is a significant difference between the methylation level in each tissue of important CpG sites for that tissue, compared to that of important CpG sites for other tissues.

### Prediction of gene interactions with the integrated public datasets

We accessed the normalized gene expression data from GeneNetwork^2^, which uniformly processed and normalized public RNA-sequencing data from 31,499 samples. To connect this expression data with our normalized public methylation data, we used the 6,067 unique eQTM genes and removed the genes that are associated with the same CpG site, which resulted in 4,052 unique eQTMs. We separately trained the CCA models for paired datasets used in eQTM discovery and for the normalized methylation data we obtained from public datasets and the public expression datasets collected from GeneNetwork.

For the paired datasets, we directly used the normalized methylation and expression values as the input data for CCA transformation. We implemented the CCA model with pyrcca^25^ and defined a Spearman correlation kernel. We finetuned the kernelized CCA with 10-fold cross validation for 1) a regularization parameter ranging from 0.001 to 1 and 2) numbers of canonical dimensions to keep ranging from 5 to 10, using “CCACrossValidate” in the Python package pyrcca^25^.

For the public dataset, we implemented the canonical correlation analysis with the PLSRegression (partial least squares regression) function in the scikit-learn Python package^24^. As a replacement for the Spearman kernel, we performed quantile normalization beforehand. As a result, we saved the first 10 components and used the default settings for other hyperparameters, namely, we used the power algorithm, a maximum of 500 iteration rounds and a tolerance value of 1×10^-5^ as the convergence criterium. As input, we provided the principal components from the public expression data as the training vectors and the components from methylation as the target vectors. We finetuned the model for the number of the input PCA components from expression and methylation data ranging from 80 to 240, with a higher coefficient of determination calculated from predicted methylation data and actual methylation data as the selection criteria. Concretely, the learned CCA takes expression profiles from individuals in the public datasets as input and can predict the corresponding methylation profiles from the same individuals. When finetuning the CCA, in each iteration, we took a new combination of PC numbers, predicted the methylation profiles, and then used the maximum coefficient of determination as the evaluation metric of this trained CCA. The coefficient of determination was calculated with the function “r2_score” from the scikit-learn Python package^24^ with “raw_values” as the setting. The best results were achieved with the CCA for negative eQTMs using 80 PCs from expression data and 240 PCs from methylation data and with the CCA for positive eQTMs using 160 PCs from the expression data and 120 PCs from methylation data.

Later, we regressed out the obtained CCA components from the public gene expression and DNA methylation datasets. With this, we obtained four different data matrices, namely, 1) the CCA components from expression data, 2) the CCA components from the DNA methylation data, 3) the residuals from the expression data and 4) the residuals matrix from the DNA methylation data. For each data matrix, we calculated pairwise correlations for each gene pair to obtain four correlation values for description for each pair. With the four correlation values, we constructed a logistic regression model with balanced loss weight to predict whether a gene pair is functionally interacting.

For training the prediction model, we took functional interaction gene pairs from STRING databases as the ground-truthing dataset. We used two different thresholds for selecting gene pairs from STRING: 400 and 600 for the “combined_score”. We used pairwise gene pairs from the eQTMs, and if a gene pair is annotated with a “combined_score” above the chosen threshold, this gene pair counted as a positive case. Otherwise, it counted as a negative case. We randomly split the dataset into training (75%) and testing datasets and trained logistic regression models with a “balanced” weight setting.

For model performance comparison, we constructed three other baseline models: 1) one with co-expression values from the public expression dataset, 2) one with co-methylation values from public methylation dataset and 3) one with both co-expression and co-methylation values. For performance evaluation, we used the AUROC and AUPRC values and the partial AUPRC values with recall value lower than 0.005. The partial AUPRC value with low recall reflects the average precision for the predictors when only considering the top few predicted gene pairs. The partial AUPRC values were obtained by calculating the area under curve values when restricting the precision values above 0.5.

### Gene enrichment analysis for CCA components

We used Gene Set Enrichment Analysis (GSEA)^26^ to find the enrichment for each CCA component identified in the public datasets. The GSEA method requires two input datasets, namely, an expression dataset and phenotype labels, and a gene set file. For the gene set file, we selected from the GSEA the file “c5.all.v7.2.symbols.gmt [Gene ontology]” containing all C5 gene sets from version 7.2 of the MSigDB database^26^. For the expression dataset, we defined the principal components as the individuals and used the principal components from each gene or each CpG site. For the phenotype labels, we defined the CCA components as the phenotypes and calculated the covariance matrix between the learned CCA components and the principal components for both expression and methylation datasets. For GSEA calculation, we selected Euclidean distance for ranking genes.

### Evaluation of predicted eQTMs with public datasets

We developed two methods to identify eQTM signals from public datasets. For both methods, we validated the method with discovered eQTMs and then applied the method to validate the predicted eQTMs. The first method calculates the average expression and methylation value per tissue from the public datasets. Then, for each CpG and gene, we correlated the resulting expression and methylation vectors. The second method uses the learned CCA transformation from the public datasets and applies the transformation to CpG sites and genes involved in discovered and predicted eQTMs. For the CCA transformation, we applied the negative and positive transformations separately. Specifically, we applied the CCA transformation learned from positive eQTMs (denoted as positive CCA transformation in the following text) to the discovered eQTMs in the test dataset and the predicted positive eQTMs and did similarly for the negative eQTMs and transformation (denoted as negative CCA transformation in the following text). For both positive and negative CCA transformations, we only used the components that were significantly associated with each other in our test dataset from the public datasets: the first four components from the negative CCA and the first three components from the positive CCA.

From both methods, we gained vector representations for each CpG site and gene of interest. Afterwards, we correlated the vectors for the discovered negative and positive eQTMs and compared the correlation distribution against the background correlation distribution. We empirically approximated the background distribution by shuffling which gene is associated with each CpG site for the known eQTMs (Supplementary Figure 16).

## Supplementary Figures Legends

**Supplementary Figure 1: The r-squared value distribution for the eQTMs in BIOS dataset.**

**Supplementary Figure 2: Comparison of the effect sizes and directions of common eQTMs for eQTMs. a** The comparison between the eQTM effect sizes discovered in this study and the Bonder, M. J. *et al*. study **b** The comparison between the eQTM effect sizes discovered in this study and the Ruiz-Arenas, C. *et al*. study.

**Supplementary Figure 3: Comparison between the effect sizes and directions of eQTM discovery results calculated from each cohort.** The dashed line in each figure indicates the FDR=0.05 calculated from each corresponding cohort.

**Supplementary Figure 4: Prediction model illustration and performance. a** The model performance of model 1, predicting whether there is an eQTM effect between the CpG site and a nearby gene. **b** The model performance of model 2, predicting the effect size direction of eQTMs.

**Supplementary Figure 5: The recall and precision values regarding the increasing threshold for probability. a** The recall and precision values regarding increasing probability threshold values for Model 1. **b** The recall and precision values regarding increasing probability threshold values for Model 2 for determining positive eQTMs. **c** The recall and precision values regarding increasing probability threshold values for Model 2 for determining negative eQTMs.

**Supplementary Figure 6: Outlier sample selection based on the first principal component for the public methylation data.** Each dot represents a sample with a unique GSM number. Red dots indicate samples with outlier PC1 values (3 standard deviation) that were discarded from further analysis.

**Supplementary Figure 7: The methylation profile of CpG sites that could be potential tissue markers.** Each value in the heatmap shows the minus log two-sided p-value from Kolmogorov-Smirnov statistic test, with the alternative hypothesis being that there is no difference between the methylation levels of the CpG markers indicated for the tissue in rows, against the methylation level of CpG markers for other tissues, in the tissue indicated in the column.

**Supplementary Figure 8: Correlation between gene–gene pairs and corresponding CpG–CpG pairs in BIOS and public datasets.** Each gray point in the scatter plots represent a gene-gene pair or a CpG site - CpG site pair. **a** Comparison between co-methylation and co-expression patterns in BIOS dataset. **b** Comparison between co-methylation and co-expression patterns in public datasets. **c** Comparison of co-methylation patterns between BIOS and public datasets. **d** Comparison of co-expression patterns between BIOS and public datasets.

**Supplementary Figure 9: Correlation between the genes *MRPL35* and *MRPS18B*. a** The co-methylation correlations of *MRPL35* and *MRPS18B* from BIOS data **b** The co-expression correlations of *MRPL35* and *MRPS18B* from BIOS data **c** The co-methylation correlations of *MRPL35* and *MRPS18B* from the public datasets **d** The co-expression correlations of *MRPL35* and *MRPS18B* from the public datasets.

**Supplementary Figure 10: Correlation between PCA components from the paired (BIOS) dataset and those from the public dataset for the eQTMs discovered from the meta-analysis. a** Correlation between PCA components from gene expression and methylation data from BIOS for negative eQTMs. **b** Correlation between PCA components from gene expression and methylation data from BIOS for positive eQTMs. **c** Correlation between PCA components from gene expression and methylation data from BIOS for negative eQTMs. **d** Correlation between PCA components from gene expression and methylation data from BIOS for negative eQTMs.

**Supplementary Figure 11: Correlation between CCA components from the paired (BIOS, ROSMAP and PIAMA) datasets for eQTMs discovered from the meta-analysis. a** Correlation between CCA components from gene expression and methylation data from BIOS for negative eQTMs. **b** Correlation between CCA components from gene expression and methylation data from BIOS for positive eQTMs. **c** Correlation between CCA components from gene expression and methylation data from BIOS for all eQTMs. This CCA transformation was trained on concatenated gene expression and DNA methylation data. **d** Correlation between CCA components from gene expression and methylation data from BIOS for all eQTMs. Before concatenating the DNA methylation data from positive and negative eQTMs, the data from negative eQTMs was timed by minus one for flipping the direction.

**Supplementary Figure 12: Correlation between CCA components from negative eQTMs, positive eQTMs, all eQTMs and all eQTMs with concatenated flipped data. a** Correlation between CCA components from gene expression and methylation data from public datasets for negative eQTMs. **b** Correlation between CCA components from gene expression and methylation data from public datasets for positive eQTMs. **c** Correlation between CCA components from gene expression and methylation data from public datasets for all eQTMs. This CCA transformation was trained on concatenated gene expression and DNA methylation data. **d** Correlation between PCA components from gene expression and methylation data from public datasets for all eQTMs. Before concatenating the DNA methylation data from positive and negative eQTMs, the data from negative eQTMs was timed by minus one for flipping the direction.

**Supplementary Figure 13: The explained variance distribution for individuals in the public datasets from negative eQTMs. a** The histograms of the explained variances of gene expression data by the each learned CCA components. **b** The histograms of the explained variances of DNA methylation data by the each learned CCA components.

**Supplementary Figure 14: Prediction comparison for gene pairs with different selection threshold from STRING database. a** The ROC, PRC and the partial PRC curves are made with a threshold of 400 for combined score in the STRING database **b** The ROC, PRC and the partial PRC curves are made with a threshold of 600 for combined score in the STRING database. The “exp” indicates the model constructed with only co-expression values. The “met” indicates the model constructed with only co-methylation values. The “exp_met” indicates the model constructed with both co-expression and co-methylation values. The “cca” indicates the model constructed with the CCA components and residuals from the proposed pipeline in this study.

**Supplementary Figure 15: Prediction scores comparison for PTM gene pairs and non-PTM gene pairs.** X-axis labels indicate the three models: “Expression” is the model built with co-expression values in public datasets only, “Expression and Methylation” is the model built with co-expression and co-methylation values in public datasets and “CCA” is the model built with CCA components and residuals from the CCA pipeline.

**Supplementary Figure 16. Correlation of gene expression and methylation levels for eQTMs in the public datasets. *a*** This illustration shows the steps for the creation of the background “Permuted CpG–gene pair” for results shown in (**b**), (**c**) and (**d**). Essentially, we calculated the pairwise Pearson correlation between all possible combinations of the CpG sites and genes identified to be significant in the meta-analysis for eQTMs and contrasted the distribution of the negative eQTMs and positive eQTMs against non-eQTMs CpG–gene combinations. **b** the results from the first method we used for summarizing the expression and methylation data for public datasets. Briefly, we took the average expression and methylation level per each tissue for the CpG sites and genes and correlated the vector of average values from tissues present in both public expression and methylation data. **c** Comparison between the CpG sites’ and genes’ CCA components correlation distribution for negative eQTMs and random CpG-gene pairs. **d** Comparison between the distribution of correlation values calculated from CCA components from negative eQTM CpG sites and negative eQTM genes, and the distribution of correlation values calculated from CCA components of random CpG-gene pairs.

**Supplementary Figure 17: The correlations between CCA components from the public dataset for predicted eQTMs. a** Comparison between the distribution of correlation values calculated from CCA components from predicted negative eQTMs and random pairs of CpG and gene. **b** Comparison between the distribution of correlation values calculated from CCA components from predicted positive eQTMs and random pairs of CpG and gene.

## Supplementary Tables Descriptions

**Supplementary Table**1. **Summary statistics for significant eQTMs from meta-analysis**

**Supplementary Table**2. **Summary statistics for significant eQTMs from BIOS**

**Supplementary Table**3. **Summary statistics for significant eQTMs from ROSMAP**

**Supplementary Table**4. **Summary statistics for significant eQTMs from PIAMA**

**Supplementary Table**5 **Predicted eQTMs**

**Supplementary Table**6. **Tissue and disease annotations for public methylation data**

**Supplementary Table**7**. Tissue composition of the public methylation data**

**Supplementary Table**8 **Tissue prediction model performance**

**Supplementary Table**9 **Metastasis prediction summary**

**Supplementary Table**10. **Metastasis predictions**

**Supplementary Table**11. **The gene–gene pairs that are highly correlated in MethylationNetwork and BIOS dataset but lowly correlated in GeneNetwork and in BIOS dataset**

**Supplementary Table**12**. Gene enrichment analysis for common components identified from public expression and methylation data.**

## Acknowledgments

We greatly acknowledge Kate Mc Intyre for her contribution to the development and refinement of texts. We greatly acknowledge Marc-Jan Bonder for sharing his scripts and experience in the DNA methylation array preprocessing steps. We greatly acknowledge dr. Chengjian Xu in Hannover Medical School for his insights and comments on the manuscript.

This project has received funding from the Netherlands Organization for Scientific Research under NWO VIDI grant number 917.164.455. L.F. is supported by grants from the Dutch Research Council (ZonMW-VIDI 917.14.374 to L.F.), by an ERC Starting Grant, grant agreement 637640 (ImmRisk), by an Oncode Senior Investigator grant and a sponsored research collaboration with Biogen.

The PIAMA study was supported by the Netherlands Organization for Health Research and Development; the Netherlands Organization for Scientific Research; the Lung Foundation of the Netherlands (with methylation studies supported by AF 4.1.14.001); the Netherlands Ministry of Spatial Planning, Housing, and the Environment; and the Netherlands Ministry of Health, Welfare, and Sport. The ROSMAP data was obtained from the AMP-AD Knowledge Portal (doi:10.7303/syn2580853). Study data were provided by the Rush Alzheimer’s Disease Center, Rush University Medical Center, Chicago. Data collection was supported through funding by NIA grants P30AG10161, R01AG15819, R01AG17917, R01AG30146, R01AG36836, U01AG32984, U01AG46152, the Illinois Department of Public Health, and the Translational Genomics Research Institute.

We thank the UMCG Genomics Coordination Center, the UMCG Research IT programme, the UG Center for Information Technology and their sponsors BBMRI-NL & TarGet for storage and compute infrastructure. We thank the Biobank-Based Integrative Omics Studies (BIOS) Consortium, funded by the Biobanking and Biomolecular Research Infrastructure Netherlands (BBMRI-NL), a research infrastructure financed by the Netherlands Organization for Scientific Research (NWO) under award number 184.021.007.

We greatly thank all the participants whose DNA methylation array data are deposited in GEO.

## Availability of data and materials

The datasets and models supporting the conclusions of this article are available in the Zenodo repository (DOI: 10.5281/zenodo.4666994) and https://data.harmjanwestra.nl/comethylation/. The code is available in GitHub (https://github.com/molgenis/methylationnetwork). The accession IDs for the public DNA methylation data can be found in Supplementary Table 6.

The DNA methylation and gene expression data from BIOS can be requested and downloaded at European Genome–Phenome Archive (EGA), accession EGAS00001001077. The data from ROSMAP can be requested and downloaded through Synapse (DOI: 10.7303/syn2580853). The PIAMA data are available upon request, and more information about its data availability can be found at https://www.birthcohorts.net/ and https://piama.iras.uu.nl/.

## Ethical compliance

All cohorts included in this study enrolled participants with informed consent, and collected and analyzed data in accordance with ethical and institutional regulations. The information about individual institutional review board approvals is available in the original publications for each cohort. Where applicable, data access agreements were signed by the investigators prior to acquisition of the data to the UMCG which state the data usage terms. To protect the privacy of the participants, data access was restricted to the investigators of this study, as defined in those data access agreements. Where applicable (ROSMAP, PIAMA, BIOS) only summary level data is made publicly available and strictly mentioned in the disclaimer that they cannot be used to re-identify study participants.

## Author Contributions

S. L. and H. W. were responsible for the cis-eQTM analysis. S. L. and H. W. were responsible for the analysis of the public DNA methylation analysis. C. Q. and G. H. K. supported the analysis with the cis-eQTM analysis with the PIAMA dataset. F. B. and P. D. supported the analysis with the GeneNetwork. N. K. processed the genotype and RNAseq data from the ROSMAP data and supported the analysis with the cis-eQTM analysis with the ROSMAP dataset. L. F. and M. S. acquired funding and supervised the study. H. W. and K. J. V. supervised S. L. in conducting the analysis. S.L. and H. W. drafted the manuscript. All authors have proof-read the final manuscript.

## Consortium

**BIOS Consortium**

Bastiaan T. Heijmans, Peter A. C. ‘t Hoen, Joyce van Meurs, Rick Jansen, Lude Franke, Dorret I. Boomsma, Jenny van Dongen, Coen D. A. Stehouwer, Cisca Wijmenga, Eline P. Slagboom, Jan H. Veldink, Hailang Mei, Maarten van Iterson, Patrick Deelen, Marc Jan Bonder, Morris A. Swertz & Wibowo Arindrarto

## Web resources

1. BIOS: https://www.bbmri.nl/services/samples-images-data/integrative-omics-data-set
2. PIAMA: https://piama.iras.uu.nl/
3. ROSMAP: https://www.synapse.org/#!Synapse:syn3219045
4. STRING: https://string-db.org/cgi/download?sessionId=bFrbbA0Zapp6
5. UniProt: https://www.uniprot.org/uniprot/?query=keyword:%22PTM%20[KW-9991]%22&fil=organism%3A%22Homo+sapiens+%28Human%29+%5B9606%5D%22+AND+reviewed%3Ayes
6. Github repository for the scripts for this project: https://github.com/molgenis/methylationnetwork
7. Zenodo repository for the data used in this project: https://zenodo.org/deposit/4666994
8. Public DNA methylation data harmonized in this study: https://data.harmjanwestra.nl/comethylation/

